# Tissue-specific increases in hypoxia-inducible factor 1 alpha protein are independent of mRNA levels during acute hypoxic exposure of the Gulf killifish, *Fundulus grandis*

**DOI:** 10.1101/2023.08.07.552360

**Authors:** Taylor E. Murphy, Jasmine C. Harris, Bernard B. Rees

## Abstract

The hypoxia inducible factor 1 (HIF1) is a central regulator of the molecular responses to low oxygen in animals. It has been extensively studied in mammals, where its tissue levels are regulated by stabilization of the alpha subunit (HIF1α) when oxygen levels decrease. Consistent with this, the initial characterization of HIF1α in fish cells in culture indicated that protein abundance increased during hypoxia even when transcription was blocked. Subsequent studies, however, have reported an increase in HIF1α mRNA levels during hypoxia in certain tissues of selected species, raising the question whether hypoxic exposure brings about coordinated changes in HIF1α mRNA and protein in tissues when measured in the same fish. We have directly addressed this question by determining levels of HIF1α protein and mRNA in the tissues of Gulf killifish, *Fundulus grandis*, exposed to short-term hypoxia (24 h at 1 mg O_2_ l^−1^). HIF1α protein was higher in brain, ovary, and skeletal muscle from fish exposed to hypoxia compared with normoxic controls by 6 h, and it remained elevated in brain and ovary at 24 h. In contrast, HIF1α mRNA levels were unaffected by hypoxia in any tissue. Moreover, levels of HIF1α protein and mRNA in the same tissues were not correlated with one another, during either normoxia or hypoxia. These results suggest that, during the initial response to low oxygen, HIF1α protein levels increase as the result of post-translational protein stabilization, rather than new transcription, as predicted from studies in mammalian and fish cells in culture.

**Summary Statement:** Parallel measurements of protein and mRNA of the hypoxia inducible factor support post-translational protein stabilization rather than new transcription in the initial response of fish to low oxygen

## Introduction

Although low dissolved oxygen (hypoxia) is a naturally occurring phenomenon in aquatic environments, its geographic scope and severity have increased in recent decades (Jenny et al., 2016; Brietburg et al., 2018). Because oxygen is critical for aerobic energy metabolism, decreased levels of oxygen have wide-ranging and frequently dramatic biological effects, including changes in behavior, impaired growth and reproduction, and increased mortality, all of which may contribute to changes in biodiversity in aquatic habitats (Vaquer-Sunyer and Duarte, 2008; Small et al., 2014; Jenny et al., 2016; Breitburg et al., 2018). When exposed to sub-lethal levels of hypoxia, many species of fish show changes in gene expression, which may improve their ability to tolerate low oxygen (Richards et al., 2009). Thus, hypoxia-tolerant fishes are valuable models for assessing the molecular responses of animals to hypoxia (Nikinmaa and Rees, 2005; Mandic et al., 2009; Richards et al., 2009), which may help determine their resilience in the current context of changing aquatic habitats.

The hypoxia-inducible transcription factors (HIFs) are evolutionarily conserved, central regulators of the molecular responses to low oxygen in animals (Majmundar et al., 2010; Semenza, 2012; Pamenter et al., 2020; Mandic et al., 2021; Townley et al., 2022). HIF1 was first identified in mammalian cell culture as a protein required for the hypoxic induction of the glycoprotein hormone, erythropoietin (Semenza et al., 1991; Maxwell et al., 1993). The active transcription factor is a dimer composed of HIF1α and HIF1β subunit, the latter of which was previously described as the aryl hydrocarbon nuclear translocator (ARNT) (Wang et al., 1995; McIntosh et al., 2010). The oxygen dependency of HIF signaling is due, in part, to post-translational regulation of HIF1α protein concentration. During normoxia, prolyl hydroxylase domain (PHD) enzymes hydroxylate HIF1α at conserved proline residues, a process that targets the alpha subunit for ubiquitin-dependent proteasomal degradation. At low oxygen levels, PHD activity decreases, HIF1α degradation is suppressed, and HIF1α protein subunits accumulate (Kaelin and Ratcliffe, 2008). HIF1α dimerizes with ARNT, translocates to the nucleus, binds to specific DNA regulatory elements in target genes, and promotes transcription (Semenza, 2012). HIF1 regulates the expression of over 100 genes, many of which improve oxygen delivery to tissues or improve the tissues’ capacity to tolerate low oxygen (Wenger et al., 2005; Ortiz-Barahona et al., 2010).

Soitamo et al. (2001) were the first to report the presence of HIF1α in fish. They sequenced HIF1α from rainbow trout (*Oncorhynchus mykiss*) and showed that HIF1α protein increased in abundance during short-term (4 h) hypoxic exposure of salmonid liver, gonad, and embryonic cells in culture. Two important observations were that the increase in HIF1α protein during hypoxia occurred when mRNA synthesis was inhibited, and that pharmacological inhibition of the proteasome during normoxia resulted in an increase in HIF1α protein. These observations suggested that HIF1α protein abundance in fish was post-translationally regulated in a similar fashion to that described in mammals. Subsequently, hypoxia-induced elevation of HIF1α protein has been reported in fish embryos, cultured cells, and adult tissues (Mandic et al., 2021 and references therein), although the extent of this elevation depends on experimental conditions (level and duration of hypoxia; temperature), species, tissue, and individual (e.g., Rissanen et al., 2006). Although Soitamo et al (2001) showed that new transcription was not required for the hypoxia-dependent increases in HIF1α protein of salmonid cells in culture, several studies have reported elevated HIF1α mRNA levels during hypoxia exposure of live fish (Mandic et al., 2021 and references therein). These studies span a range of taxa, tissues, and experimental conditions, and when considered in aggregate, HIF1α mRNA levels were elevated in about half of the samples from hypoxia-exposed fish (Dataset S1). When data were analyzed by tissue, hypoxia-dependent increases in HIF1α mRNA were most frequent for kidney, followed by liver, gill, and brain, and least frequent in heart and skeletal muscle (Fig. 1). In addition to this apparent tissue dependence, HIF1α mRNA was more likely to be elevated in fish exposed to lower oxygen levels, although the time course of expression was highly variable (Dataset S1).

**Fig. 1.**
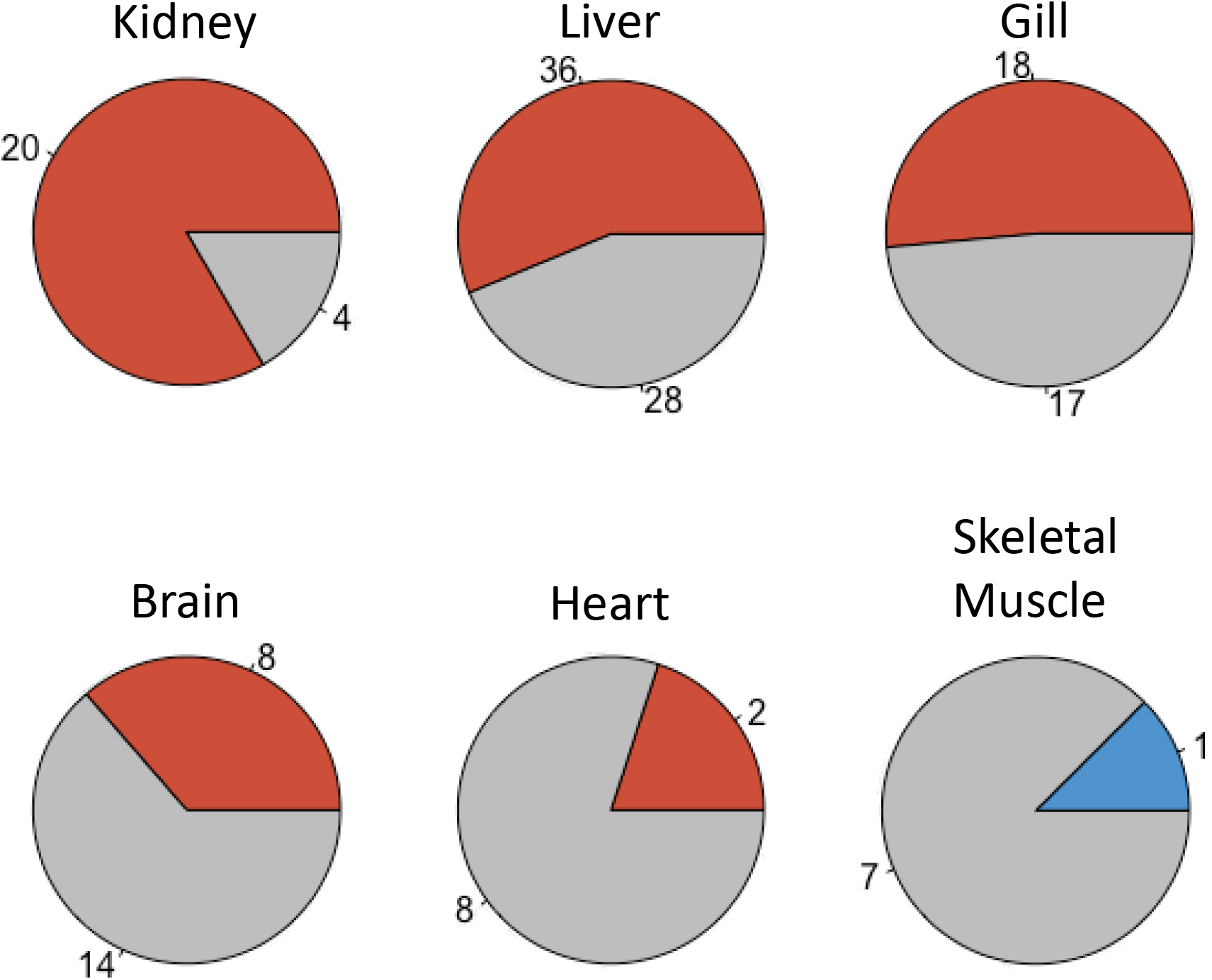
Summary of the effects of hypoxic exposure on HIF1α mRNA in tissues of fish. Data for HIF1α mRNA abundance determined by qPCR were extracted from 15 studies published between 2001 and 2023. Fish were exposed to dissolved oxygen concentrations ranging from 0.7 to 4.5 mg O_2_ l^−1^ and tissues were harvested after 0.5 to 300 h. The results are displayed as pie charts by tissue: A, brain; B, gill; C, heart; D, kidney; E, liver; F, skeletal muscle. Colors indicate the proportion of all tissue samples in which HIF1α mRNA abundance was greater in hypoxia than in normoxia (red); HIF1α mRNA was lower during hypoxia than normoxia (blue); HIF1α mRNA was not significantly different between hypoxia and normoxia (grey). The numbers on the edges of the pie charts indicate the number of samples having increased, decreased, or unchanged HIF1α mRNA during hypoxia. Note, each sample reported in each study (tissue, duration of hypoxia, level of hypoxia) was counted as a single sample in this analysis, resulting in studies having disproportionate weights. For more details, see Dataset S1.

The myriad experimental conditions among previous studies makes it difficult to ascertain whether changes in the abundance of HIF1α mRNA and protein are linked. For example, if HIF1α mRNA is elevated, does this precede or occur contemporaneously with elevated HIF1α protein, e.g., suggestive of transcriptional control of HIF1α protein abundance? Or, does HIF1α protein increase prior to changes in HIF1α mRNA, which could indicate positive feedback regulation? In the limited number of studies that report both HIF1α mRNA and protein, their levels are either weakly associated or unrelated (Soitamo et al., 2001; Law et al., 2006; Rissanen et al., 2006; Sollid et al., 2006; Robertson et al., 2014; Guan et al., 2017). Importantly, none of these earlier studies compared tissue levels of HIF1α protein and mRNA to one another among individuals.

In this study, therefore, we examined the relationship of HIF1α mRNA and protein in tissues of the Gulf killifish *Fundulus grandis* Baird & Girard 1853 exposed to acute hypoxia. *Fundulus grandis* is a widespread and ecologically important species of the salt marsh communities of the Gulf of Mexico (Nordlie, 2006), areas that are prone to natural and anthropogenic hypoxia (Engle et al., 1999). Previous research showed that HIF1α protein increased in several tissues of *F. grandis* exposed to 1 mg O_2_ l^−1^ (∼13% of the air-saturated oxygen concentration) for 24 h (Gonzales-Rosario, 2016). Here, we asked whether this increase in HIF1α protein was associated with an increase in HIF1α mRNA levels when measured in the same tissues among individuals. Fish were subjected to hypoxia and their tissue levels of HIF1α protein were determined by immunoprecipitation and western blotting and levels of HIF1α mRNA were measured by quantitative PCR. The results show that HIF1α protein accumulation occurs in a tissue- and individual-dependent fashion that is independent of changes in HIF1α mRNA. In addition, because HIF1α is part of a larger gene family (Rytokonen et al., 2011; Graham and Presnell, 2017; Townley et al., 2022), we took the opportunity to measure the mRNA abundance of HIF2α and HIF3α. Similar to HIF1α, we found no evidence for changes in these transcripts in tissues of *F. grandis* subjected to acute hypoxia. Thus, we conclude that the initial increase in HIF1α protein in tissues of acutely exposed *F. grandis* is due to post-translational mechanisms as in mammals (Kaelin and Ratcliffe, 2008) and salmonid cells in culture (Soitamo et al., 2001).

## Materials and Methods

### Fish collection and maintenance

Adult female *F. grandis* (n = 27; mean mass, 16.29 g; range, 10.17-24.93 g) were purchased in February 2019 and housed at the University of New Orleans. Fish were treated within one week for bacterial infection and ectoparasites with API Furan-2 and API General Cure (Chalfont, PA, USA). Fish were housed in two 38-l aquaria in dechlorinated water that was adjusted to a salinity of 9 – 12 using Instant Ocean Synthetic Sea Salt (Blacksburg, VA, USA). Water was maintained at 24.6 ± 1.2°C (mean ± range), aerated to maintain > 80% (87.3 ± 4.4%) air saturation, and filtered. Dissolved oxygen (DO, in % saturation and mg O_2_ l^−1^), temperature, and salinity were monitored daily using a YSI Pro2030 oxygen-temperature-salinity probe (Yellow Springs, OH, USA). Nitrates, nitrites, and ammonia were measured once a week, and partial water changes (∼25%) were conducted as needed to keep these variables within acceptable levels. Fish were fed 1-1.5% of their body mass daily between 10:00 and 12:00 with TetraMin Tropical Flake fish food (Blacksburg, VA, USA) and maintained under a 12 h light: 12 h dark photoperiod. Procedures for handling and experiments adhered to established guidelines approved by the University of New Orleans Institutional Animal Care and Use Committee (University of New Orleans IACUC Protocol 18-006).

### Experimental exposures

Experimental exposures were conducted between one and two months after collection. The day before experiments, fish were fed in the morning (22 to 28 h before exposures), transferred to the exposure tank in the evening, and allowed to adjust to the tank overnight. The 76-l exposure tank was subdivided by polystyrene grate into four individual fish compartments, each of which had a removable a polystyrene grate below the air-water interface to prevent fish from accessing the surface. The four compartments were separated by opaque dividers, preventing fish from seeing one another. Water in the exposure tank was thoroughly circulated throughout the tank by submersible water pumps, and it had the same composition as the maintenance tanks except for dissolved oxygen (see below).

Groups of two to four fish were randomly selected from the holding tanks and assigned to one of four treatments: 6 h normoxia; 24 h normoxia; 6 h hypoxia; 24 h hypoxia. These treatments were repeated with independent groups of fish to achieve the final sample sizes: 6 h normoxia, n = 6; 24 h normoxia, n = 4; 6 h hypoxia, n = 9; 24 h hypoxia, n = 8. For normoxic exposures, water was continuously aerated with room air to maintain > 7 mg l^−1^ DO (∼90% air-saturation). For hypoxic exposures, nitrogen gas was bubbled into the exposure tank under the control of a CanaKit Raspberry Pi 3 Model B+ (North Vancouver, BC, Canada), which received input from a galvanic oxygen electrode (Atlas Scientific, Long Island City, NY, USA). Nitrogen was introduced to achieve the target DO of 1 mg O_2_ l^−1^ (∼13% air-saturation), which took 45 – 60 min, and then as needed to maintain the target. The exposure tank was continuously bubbled with air at a low rate to prevent the DO from dropping below the target. Plastic bubble wrap was placed on the surface of the water to minimize diffusion of oxygen from ambient air. Manual measurements with the YSI Pro2030 demonstrated that oxygen, temperature, and salinity were uniform throughout the exposure tank.

### Euthanasia and tissue sampling

After 6 or 24 h exposure to either normoxia or hypoxia, fish were netted and immersed in a slurry of aquarium water and ice (< 2°C) until loss of equilibrium (Larter and Rees, 2017). Fish were briefly blotted and bled by severing the caudal peduncle. Blood samples (30-50 µl) were collected in heparinized capillary tubes and analyzed for indicators of hypoxic exposure (see below). Euthanasia was confirmed by severing the spinal cord behind the head, after which, fish were weighed and dissected for brain, ovaries, skeletal muscle, liver, and gills. Tissues were frozen in liquid nitrogen, placed on dry ice, and transferred within 2 h to −80°C, where they were kept until analysis (within 6 months). Tissue sampling occurred from 13:30 to 16:30.

### Blood variables

For each fish, hematocrit was determined by centrifugation of one blood sample in a microhematocrit centrifuge (BD Clay Adams AutoCrit Ultra 3, Franklin Lakes, NJ, USA) for 3 min. A second blood sample was collected and immediately diluted into 1.0 ml of saline consisting of 145 mM NaCl, 5 mM KCl, 12 mM NaHCO_3_, 3 mM NaH_2_PO_4_, with 3 mM sodium citrate (dihydrate) to prevent clotting (final pH 7.6) (Genz and Grosell, 2011; Wood *et al*., 2010). Red blood cells (RBC) were counted in the diluted blood using a Neubauer hemocytometer. The same diluted blood sample was frozen at −20°C, thawed, and vortexed vigorously to lyse red blood cells. Lysates were centrifuged at 16,000 x g for 60 s, and hemoglobin concentration was determined in the supernatant with a 96-well plate hemoglobin assay (Cayman Hemoglobin Colorimetric Assay Kit, Ann Arbor, MI, USA). RBC count (cells per ml), blood hemoglobin (Hb; g dl^−1^), and mean corpuscular hemoglobin concentration (MCHC; g Hb l^−1^ cells) were determined after accounting for the dilution of whole blood by saline.

A third blood sample was collected and prepared for blood glucose and lactate measurements as described by Larter and Rees (2017). Glucose was determined using a glucose oxidase-peroxidase coupled colorimetric assay according to the manufacturer’s directions (Glucose Colorimetric Assay Kit, Ann Arbor, MI, USA). Lactate was determined as described by Virani and Rees (2000) with the following modifications for a 96-well plate format. The final assay volume was 250 µl, and contained glycine (192 mM), hydrazine (160 mM), NAD^+^ (5 mM), LDH (12 units/ml), and lactate standard or sample. Each standard and sample was assayed in quadruplicate, with an equal volume of water replacing LDH in two wells to account for non-specific absorbance. Plates were agitated for 5 s, incubated at 37°C for 1 h, and read at 340 nm using a Molecular Devices Versamax plate reader (San Jose, CA, USA).

### Tissue lysate preparation and HIF1#x03B1; protein analysis

Frozen tissues were rapidly weighed, and 50-100 mg was ground under liquid nitrogen in pre-cooled mortars and pestles. Frozen tissue powder was added to 1 ml lysis buffer [137 mM NaCl, 2.7 mM KCl, 10 mM Na2HPO4, 1.8 mM KH2PO4, containing 1% Igepal, 0.5% sodium deoxycholate, 0.1% SDS, 1mM EDTA, 1mM sodium orthovanadate, 50 µg MG-132 ml^−1^ (Fisher Scientific, Lenexa, KS, USA) and 1% protease inhibitor cocktail (Sigma-Aldrich, St. Louis, MO, USA)] and homogenized with 2 x 10 strokes using Teflon-glass homogenizers (Thomas Scientific, Houston, TX, USA) on ice. Lysates were centrifuged at 4°C for 10 min at 10,000 x g, and the supernatants were frozen at −80°C. Protein concentration was determined by the bicinchoninic acid protein assay (Smith *et al*., 1985) using bovine serum albumin as the standard (Pierce, ThermoFisher, Rockford, IL, USA).

HIF1α was immunoprecipitated from tissue lysates as described in Gonzales-Rosario (2016) using chicken polyclonal antibodies developed against HIF1α from *F. heteroclitus* (Townley *et al*., 2017). All steps were conducted at 0-4°C unless otherwise indicated. In brief, a volume of lysate containing 1-2 mg protein (1 mg in brain; 1.25 mg in skeletal muscle; 2 mg in ovaries, liver, and gills) was brought to a final volume of 1.0 ml with lysis buffer. Reactions were cleared of non-specific binding by incubating with 25 µl of PrecipHen reagent (agarose-coupled goat anti-chicken IgY; Aves Lab; Davis, CA, USA) for 30 min with gentle rocking. Reactions were centrifuged at 1500 x g for 10 min. The supernatants were removed to clean tubes and incubated with affinity-purified HIF1α antibody (5 µl, equivalent to 5 µg IgY) for 1 h, after which PrecipHen reagent (25 µl) was added to each reaction and incubated with end-over-end rocking overnight (16 – 20 h). Precipitated antigen-antibody-PrecipHen complexes were washed sequentially in 0.5 ml each of TBS (20mM Tris, pH 7.6, 150 mM NaCl) containing 0.05% Tween-20, TBS, and 50 mM Tris pH 6.8, centrifuging each time at 1,500 *g* for 5 min. The final immunoprecipitates were resuspended in Laemmli sample buffer (Laemmli, 1970) containing 50 mM dithiothreitol, heated at 95°C for 3 minutes, clarified by centrifugation, and stored at −20°C.

Immunoprecipitate proteins were separated by sodium dodecyl sulfate polyacrylamide gel electrophoresis at 200 V for 50 – 60 minutes (Laemmli, 1970). Gels included molecular weight markers (Cell Signaling Biotinylated Protein Ladder, Danvers, MA, USA; Novex Sharp Pre-stained Protein Standard, Waltham, MA, USA) and positive HIF1α controls (in vitro transcribed and translated *F. heteroclitus* HIF1α; Townley *et al*., 2017). After electrophoresis, proteins were transferred to polyvinylidene difluoride membranes by electrotransfer at 100 V for 1 h at 10°C in 25 mM Tris, 192 mM glycine, 20% methanol, 0.05% SDS (Towbin *et al*., 1979). The efficiency of electrotransfer was verified by the absence of proteins in gels stained by colloidal Coomassie blue (Neuhoff et al., 1988). Blots were blocked in TBS containing 0.05% Tween-20 (TBS-T) and 5% non-fat dry milk at room temperature for 1 h, followed by incubation in the same buffer containing 1:500 dilution of chicken anti-HIF1α. After overnight incubation in primary antibody at 4°C, blots were washed with TBS-T three times and then incubated in TBS-T containing 5% non-fat dry milk and anti-biotin horseradish peroxidase (Cell Signaling, Danvers, MA, USA) and HRP-conjugated donkey anti-chicken antibody (Sigma-Aldrich, St. Louis, MO, USA) diluted to 1:2000 and 1:5000, respectively. Blots were washed with TBS-T five times, developed in enhanced chemiluminescent detection reagents (Harlow and Lane, 1988) at room temperature for 60 s, and imaged with a ChemiDoc MP (Bio-Rad, Hercules, CA, USA). The resulting images were analyzed with ImageLab (Bio-Rad) after automatic background subtraction.

### mRNA preparation and analysis

Frozen samples of skeletal muscle, liver, ovary, and gill (approximately 15-20 mg each) were ground under liquid nitrogen in pre-cooled mortars and pestles, and extracted for RNA using commercial kits (RNeasy fibrous tissue kit, Qiagen, Valencia, CA). The manufacturer-supplied protocol was modified to include a second round of DNase I treatment in solution to ensure no genomic contamination. The concentration of RNA was measured using a NanoDrop ND-1000 spectrophotometer (NanoDrop Technologies, Wilmington, DE). All samples had A260:A280 ratios between 1.8 and 2.2, indicating high RNA purity. RNA integrity was determined using a 2100 Bioanalyzer (Agilent, Santa Clara, CA), and only samples with RNA integrity numbers greater than 7.3 were used in subsequent steps. Total RNA was diluted to final concentrations of 500 ng µl^−1^ for ovary and muscle samples or 1 µg µl^−1^ for liver and gill, and RNA contained in 1 µl was reverse transcribed by TaqMan reverse transcriptase with random hexamer primers (Applied Biosystems, Foster City, CA). Samples lacking the reverse transcriptase were prepared simultaneously as controls to ensure the absence of genomic DNA.

Quantitative real time PCR (qPCR) for HIF1α, HIF2α, HIF3α, and ARNT was carried out using an iQ5 real-time PCR detection system (Bio-Rad) and gene-specific primers (Townley et al. (2017). Each qPCR reaction was performed in 25 µl reactions which contained 12.5 µl SYBR Green PCR Master Mix (Applied Biosystems), 0.1 µM of each primer, and 1µl of cDNA. The qPCR protocol followed a denaturation step for 20 s at 90°C, then 40 cycles of 95°C for 10 s and 60°C for 30 s. Melting curves were obtained by increasing temperature from 50 to 95°C in 90 steps of 0.5°C to ensure the presence of a single amplicon. All qPCR assays were performed in duplicate except when the standard deviation between the duplicates was more than 0.5 cycle, in which case, assays were done in quadruplicate. A pooled cDNA sample was formed by combining equal volumes of cDNA from at least six individuals per treatment group. This pooled sample was serially diluted to assess primer efficiencies, which ranged from 90 – 105%. The cDNA pooled sample was also included in every 96-well plate for a given gene in order to standardize run-to-run variation. Finally, several randomly selected samples without reverse transcriptase treatment confirmed that genomic DNA was not amplified.

Relative mRNA abundance was computed according to Pfaffl (2001), which calculates mRNA levels relative to a control treatment and levels of a control gene, after accounting for the amplification efficiencies for different genes. For this calculation, the average threshold cycle number of all normoxic samples (6 and 24 h) for a given gene was used as the control against which each sample for that gene was compared. To control for variation in RNA among samples, the threshold cycle number for HIFα was normalized by the threshold cycle number for ARNT, the HIF1α beta subunit, which is not affected by hypoxic treatment (Majmundar et al., 2010).

### Statistical analyses

Normality of response variables was determined using Shapiro-Wilk tests, and equality of group variances was tested using Bartlett’s tests. Variables that were not normally distributed were transformed using the Box-Cox technique (Box and Cox, 1964). The effects of hypoxia treatment, exposure time, and the interaction between treatment and time were assessed by two-way analysis of variance (ANOVA). Relationships between selected response variables were assessed by Spearman rank order correlation, and false-discovery rate corrections were performed using the R package (p.adjust). Statistical analyses were performed in R version 3.6.1 (R Studio Team, 2020) and graphs were created in GraphPad Prism version 7.01 (GraphPad Software, Boston, MA, USA).

## Results

### Indicators of hypoxia exposure

Blood variables reflecting oxygen transport capacity and carbohydrate metabolism were measured in *F. grandis* to verify that the experimental exposures caused oxygen limitation (Fig. 2). Exposure of fish to ∼1 mg O_2_ l^−1^ resulted in an increase in hematocrit (Fig. 2A), as expected. Although there was a trend toward increased red blood cell count, this change was not significant (Fig. 2B), suggesting that the major determinant of higher hematocrit during acute hypoxia was an increase in mean corpuscular volume (MCV). Indeed, MCV calculated for hypoxic fish (83 × 10^−15^ l at 6 h and 93 × 10^−15^ l at 24 h) was significantly greater (2-way ANOVA, p_Trx_ = 0.015) than that for normoxic fish (70 × 10^−15^ l at 6 h and 77 × 10^−15^ l at 24 h). Consistent with the lack of an effect of hypoxia on RBC count, there was no significant difference in blood hemoglobin (Hb) concentration between normoxic and hypoxic fish (Fig. 2C). Because hypoxia led to an increase in MCV with no increase in blood Hb, hypoxia caused a significant decrease in mean corpuscular hemoglobin concentration (MCHC) (Fig. 2D). Hypoxia also led to moderate hyperglycemia (Fig. 2E) and a significant increase in blood lactate (Fig. 2F). There was no effect of time of exposure, nor interactions between hypoxia treatment and exposure time, for any variable, indicating that these physiological responses to hypoxia occurred within the first 6 h of exposure.

**Fig. 2.**
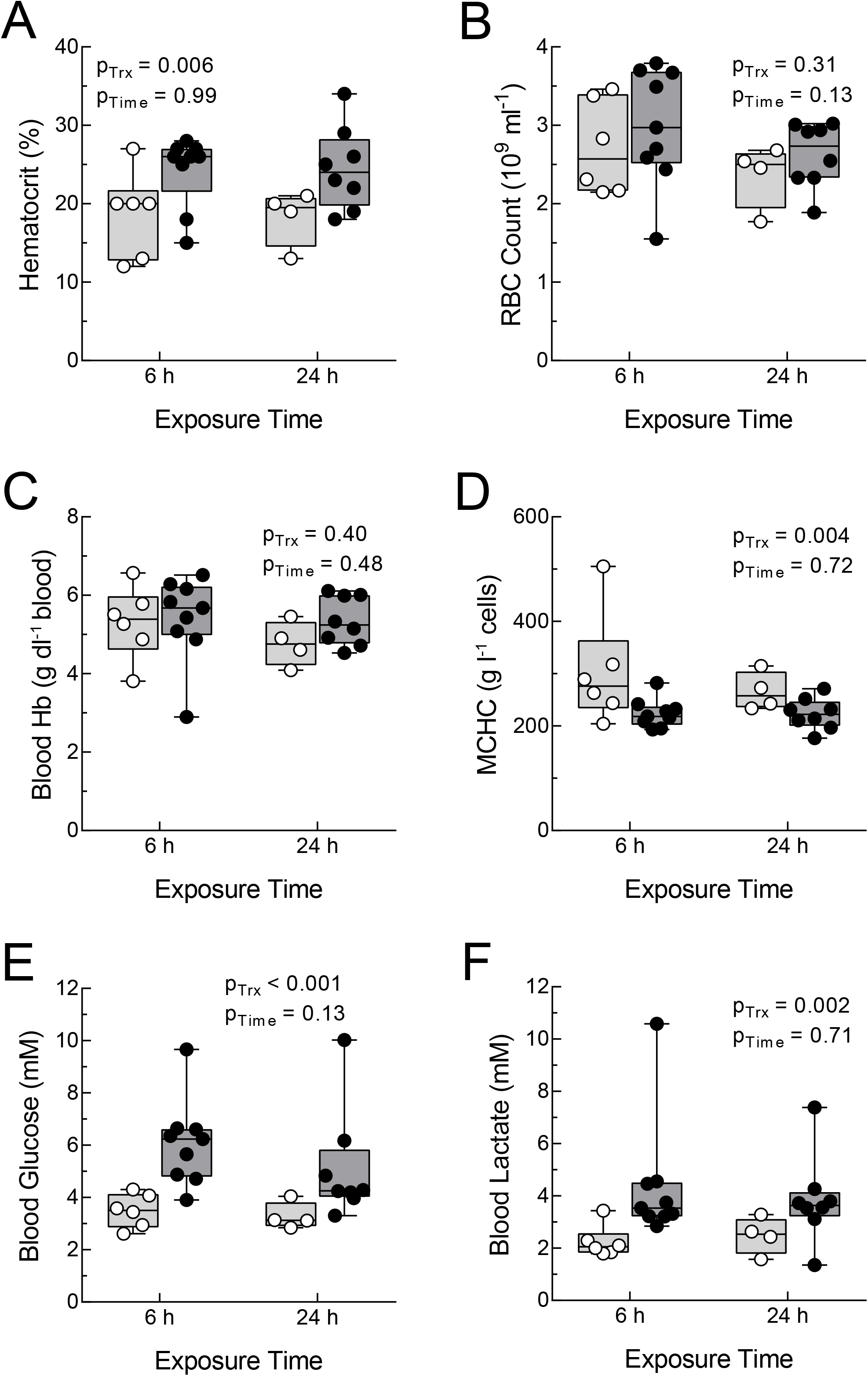
Blood indicators of oxygen transport and carbohydrate metabolism in *Fundulus grandis* during acute hypoxia. Fish were exposed to normoxia (> 7 mg O_2_ l^−1^; light grey boxes and open symbols) or hypoxia (∼1 mg O_2_ l^−1^; dark grey boxes and closed symbols) for 6 or 24 h and sampled for hematocrit (A), red blood cell (RBC) count (B), blood hemoglobin (Hb) (C), mean corpuscular hemoglobin concentration (MCHC) (D), blood glucose (E), and blood lactate (F). Lines of box and whiskers plots represent (from lowest to highest) the minimum, the 25^th^ percentile, the median, the 75^th^ percentile, and the maximum. The effects of hypoxia treatment (p_Trx_), exposure time (p_Time_), and the interaction between treatment and time were determined by 2-way ANOVA (all interactions were non-significant and not shown).

### HIF1α protein abundance

Levels of HIF1α protein, determined by immunoprecipitation followed by western blotting, increased in *F. grandis* exposed to hypoxia in a tissue- and individual-dependent fashion (Fig. 3). An image of a western blot of HIF1α protein immunoprecipitated from brain (Fig. 3A) shows that most of the hypoxic samples (lanes 3-4, 6-7, and 9) had higher levels of HIF1α protein than normoxic samples (lanes 2, 5, 8, and 10-11), although there was appreciable variability in HIF1α protein abundance in both hypoxic and normoxic fish. Some of this variation could be due to differences in the effectiveness of immunoprecipitation among samples. Thus, to account for immunoprecipitation efficiency, the relative HIF1α protein abundance for each sample was determined by dividing the HIF1α band intensity by the intensity of the immunoprecipitating IgY band for the same sample.

**Fig. 3.**
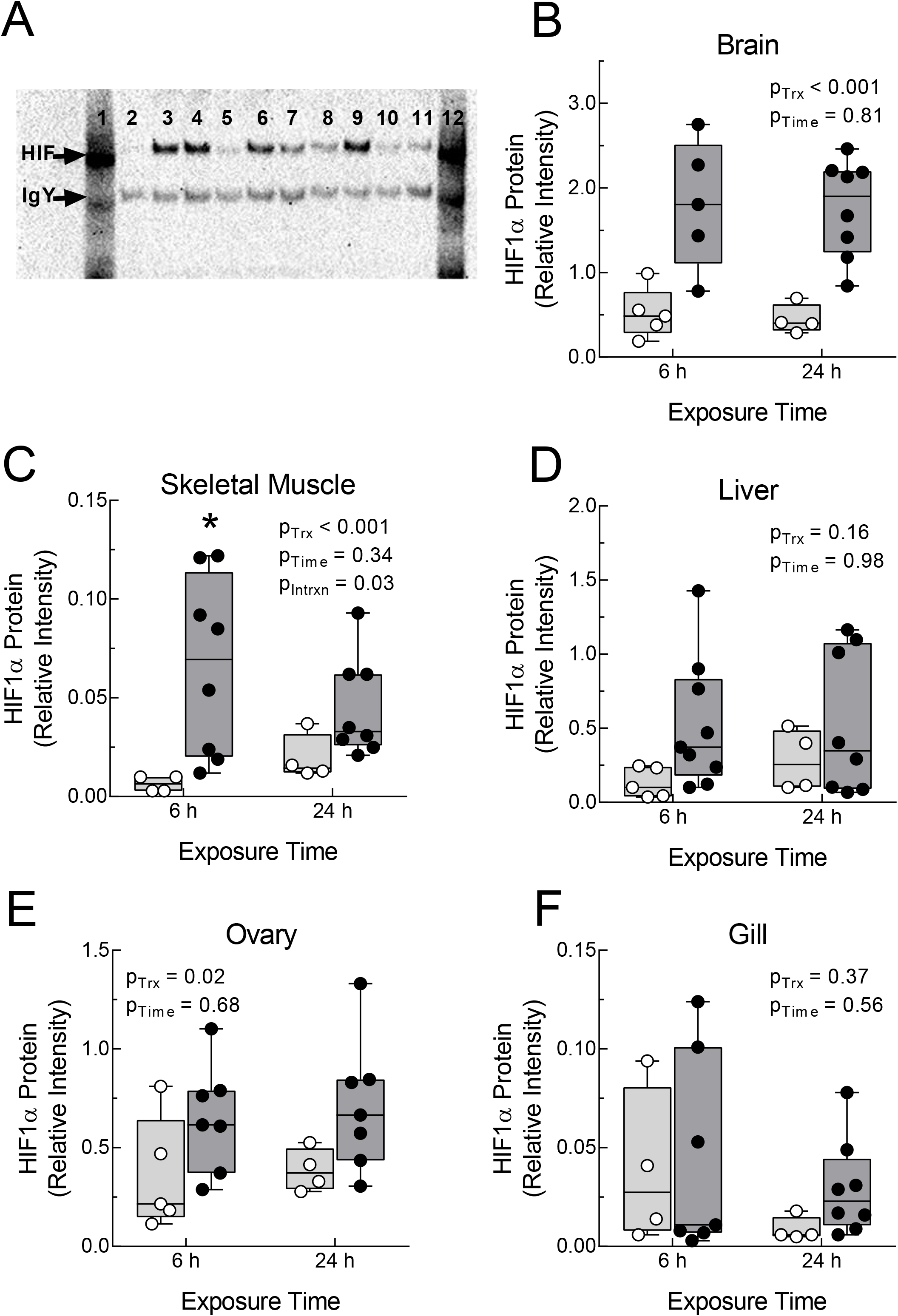
HIF1α protein levels in tissues of *Fundulus grandis* during acute hypoxia. Panel A shows a western blot of HIF1α immunoprecipitated from brain lysates of fish exposed for 6 h to normoxia (> 7 mg O_2_ l^−1^; lanes 2, 5, 8, 10, and 11) or hypoxia (∼1 mg O_2_ l^−1^; lanes 3-4, 6-7, and 9). A positive HIF1α control is shown in lane 1 and the mobility of HIF1α and chicken IgY are shown by arrows (at left). HIF1α protein abundance in each sample was expressed as the ratio of the HIF1α band intensity to the IgY band intensity. Relative HIF1α protein abundance was determined for brain (B), skeletal muscle (C), liver (D), ovary (E), and gill (F) after exposure of fish to normoxia (> 7 mg O_2_ l^−1^; open symbols and light boxes) or hypoxia (∼1 mg O_2_ l^−1^; filled symbols and dark grey boxes) for 6 or 24 h. Lines of box and whiskers plots represent (from lowest to highest) the minimum, the 25^th^ percentile, the median, the 75^th^ percentile, and the maximum. The effects of hypoxia treatment (p_Trx_), exposure time (p_Time_), and the interaction between treatment and time were determined by 2-way ANOVA. The treatment by time interaction was only significant for skeletal muscle, for which the effect of hypoxia on HIF1α abundance was significant at 6 h exposure (asterisk).

The effects of hypoxia on relative HIF1α protein abundance were significant in brain (Fig. 3B) and ovary (Fig. 3E). In brain, HIF1α protein levels were about 3-times higher in hypoxia than in normoxia, and in ovaries, HIF1α protein levels were about 2-fold higher in hypoxia. In these two tissues, there was no effect of exposure time nor was there an interaction between hypoxia treatment and exposure time. In skeletal muscle, there was a significant interaction between treatment and exposure time, with significantly elevated HIF1α protein levels in hypoxic fish at 6 h compared to corresponding controls (Fig. 3C). For liver (Fig. 3D) and gill (Fig. 3F), there were no significant effects of treatment, exposure time, or their interaction. Of note, levels of HIF1α protein determined by immunoprecipitation were highest in brain, followed by ovary and liver. HIF1α protein levels were extremely low in skeletal muscle and gill during both normoxia and hypoxia.

Correlation analyses were conducted to assess whether HIF1α protein levels were consistently low or high across tissues among individuals within treatment groups (Supplemental Table S1). After correction for multiple comparisons, tissue levels of HIF1α protein were not correlated among tissues of either normoxic or hypoxic fish.

### HIF1α mRNA abundance

Levels of HIF1α mRNA were determined by qPCR in four tissues (the small size of brain tissue required that it be used exclusively for HIF1α protein determination). HIF1α mRNA levels were normalized by the levels of ARNT mRNA, the transcript for the dimeric partner of HIF1α, which is not affected by hypoxic treatment (Majmundar et al., 2010). Thus normalized, there were no significant effects of hypoxia treatment, exposure time, or their interaction on HIF1α mRNA levels in any tissue (Fig. 4). The only change that approached statistical significance was a trend toward lower levels of HIF1α mRNA in skeletal muscle during hypoxia (p_Trx_ = 0.07; Fig. 4A). As observed for HIF1α protein levels, correlation analyses showed that tissue levels of HIF1α mRNA were not significantly related across tissues, in either normoxia or hypoxia (Supplemental Table S2).

**Fig. 4.**
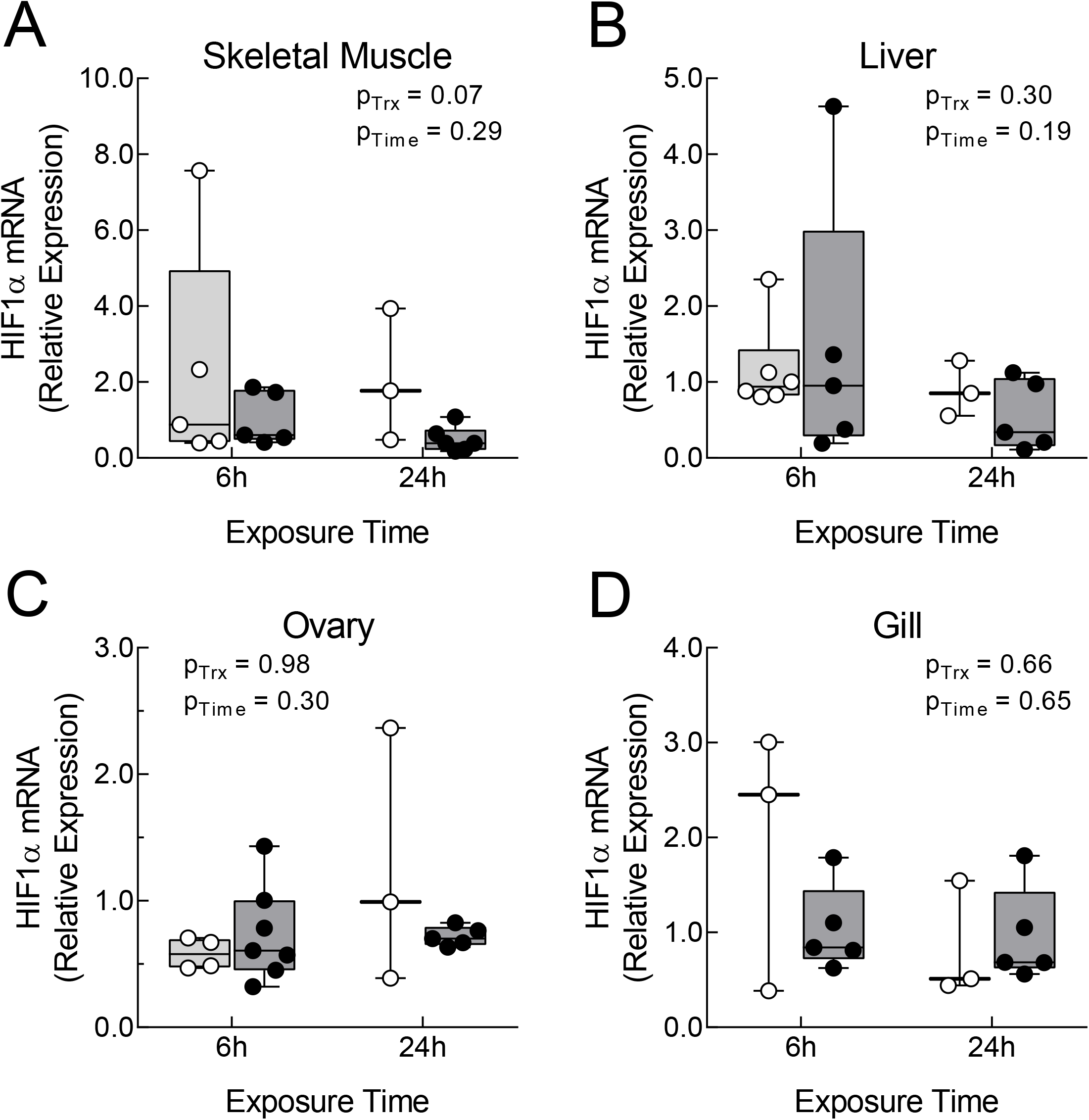
HIF1α mRNA levels in tissues of *Fundulus grandis* during acute hypoxia. Levels of HIF1α mRNA were determined by qPCR and expressed relative to the level of ARNT mRNA in the same samples. Relative HIF1α mRNA abundance was determined for skeletal muscle (A), liver (B), ovary (C), and gill (D) after exposure of fish to normoxia (> 7 mg O_2_ l^−1^; open symbols and light grey boxes) or hypoxia (∼1 mg O_2_ l^−1^; filled symbols and dark grey boxes) for 6 or 24 h. Lines of box and whiskers plots represent (from lowest to highest) the minimum, the 25^th^ percentile, the median, the 75^th^ percentile, and the maximum. The effects of hypoxia treatment (p_Trx_), exposure time (p_Time_), and the interaction between treatment and time were determined by 2-way ANOVA (all interactions were non-significant and not shown).

The copy number of HIF1α mRNA transcripts were determined from dilutions of known concentrations of a plasmid encoding HIF1α (Townley et al., 2017). When analyzed by 2-way ANOVA, both the effects of hypoxia treatment and exposure time on gill HIF1α copy number approached statistical significance (0.10 > p > 0.05) (Supplemental Table S3). However, these trends were mirrored by higher ARNT in the same samples, suggesting that this effect was not specific to HIF1α (Supplemental Table S3). The levels of HIF2α and HIF3α mRNA were also determined in the same tissues. There were no effects of hypoxia treatment on HIF2α or HIF3α mRNA levels, either expressed relative to ARNT mRNA levels (Supplemental Figs. S1 and S2) or as copy number (Supplemental Table S3). Of note, the absolute level of HIF2α was the highest of all HIFα transcripts and particularly elevated in gill tissue (Supplemental Table S3), as documented previously in *F. heteroclitus* and other fishes (Townley et al., 2017; 2022)

### Correlation between HIF1α protein and mRNA

The observations that HIF1α protein changes in certain tissues (Fig. 3) and HIF1α mRNA does not (Fig. 4) strongly argue that the levels of these two macromolecules are not linked during acute hypoxic exposure of *F. grandis*. Given that levels of HIF1α protein and mRNA both displayed considerable variation (Figs. 3, 4); however, it was possible that their abundances covary among individuals within treatment groups, even if there were no apparent effects of hypoxia on the mean values. Thus, the correlations between tissue levels of HIF1α protein and mRNA among individuals were examined in both treatment groups (Fig. 5). In none of the tissues examined was the correlation between HIF1α protein and HIF1α mRNA significant, during either normoxia or hypoxia (Table 1).

**Fig. 5.**
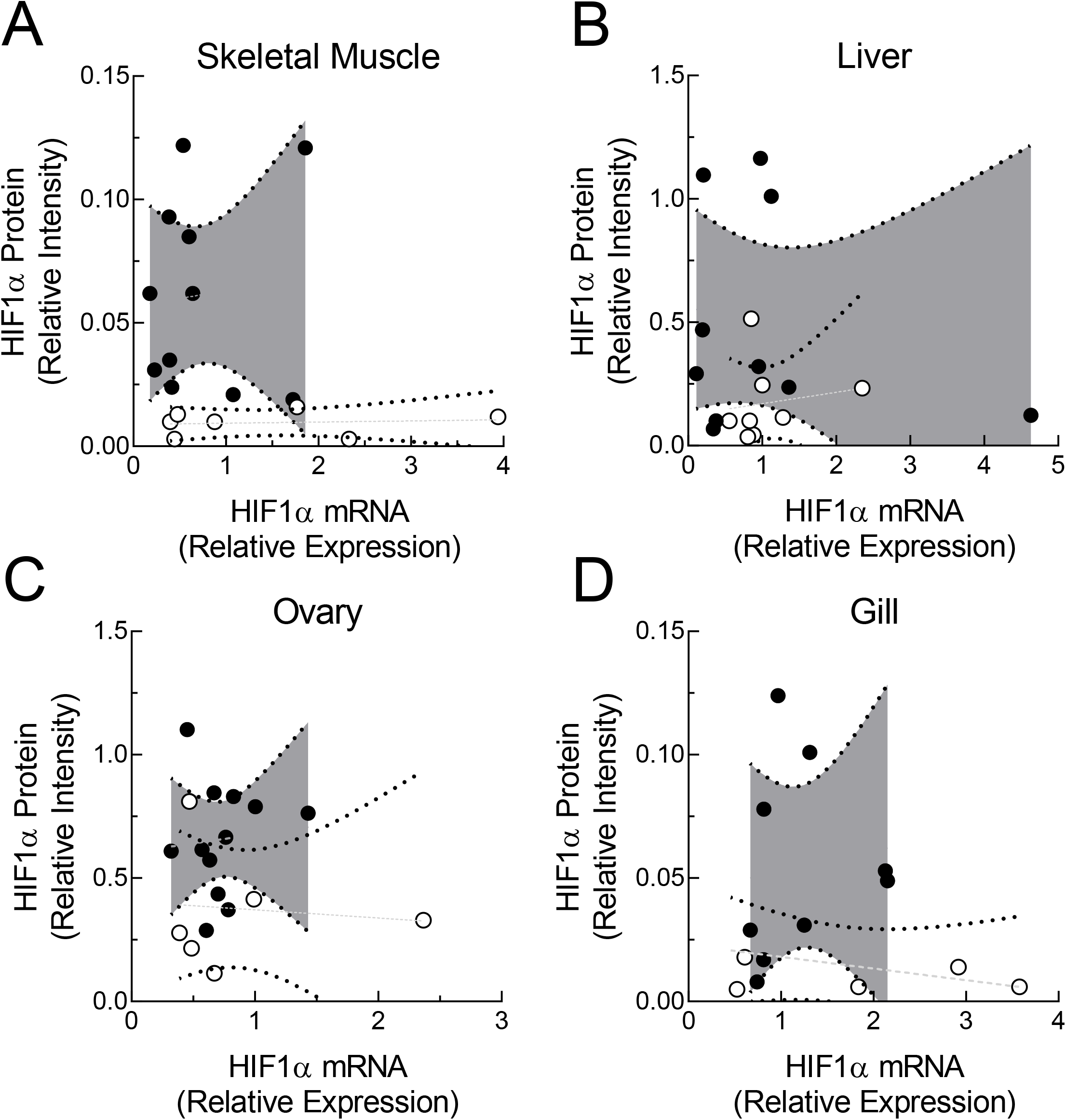
The relationship between HIF1α protein and mRNA levels in tissues of *Fundulus grandis* during acute hypoxia. The relative abundance of HIF1α protein and HIF1α mRNA were determined in skeletal muscle (A), liver (B), ovary (C), and gill (D) and plotted against one another pooling time points (6 and 24 h) within treatment groups. Individual samples are shown as points and 95% confidence bands were determined by least squares linear regression of normoxic samples (open symbols and light grey confidence bands) and hypoxic samples (filled symbols and dark grey confidence bands).

**Table 1.** Correlation analyses of HIF1α protein and HIF1α mRNA levels in *Fundulus grandis* tissues during normoxia or after acute exposure to hypoxia.

| Tissue | Normoxia |  | Hypoxia |  |
| --- | --- | --- | --- | --- |
|  | rho | P | rho | P |
| Muscle | 0.09 | 0.85 | -0.18 | 0.64 |
| Liver | 0.50 | 0.20 | -0.28 | 0.40 |
| Ovary | -0.40 | 0.50 | 0.18 | 0.63 |
| Gill | -0.32 | 0.54 | 0.53 | 0.14 |

Note: For each sample, HIF1α protein was expressed as the band intensity in western blots relative to the intensity of IgY band in the same sample, and HIF1α mRNA was expressed relative to ARNT mRNA measured in the same sample (see Methods). Correlations were assessed by Spearman's rank order correlation coefficient (rho). False-discovery rate-adjusted probabilities (P) are shown. Separate analyses were done for normoxic and hypoxic samples, pooling 6 h and 24 h timepoints within each treatment.

## Discussion

### Validation of experimental exposure

The main goal of this study was to determine the levels of HIF1α mRNA and protein in tissues of the Gulf killifish, *F. grandis*, when exposed to short-term hypoxia (6 or 24 h at 1 mg O_2_ l^−1^ or ∼13% of air-saturation at 25°C). These exposure conditions led to higher hematocrit, blood glucose, and blood lactate, all of which are typical responses of fish to low oxygen availability (Gallaugher and Farrell, 1998; Virani and Rees, 2000; Larter and Rees, 2017; Richards, 2009). The increase in hematocrit was mainly accounted for by RBC swelling, rather than an increase in RBC number. This increase in RBC volume was likely due to adrenergic stimulation of erythrocytic Na^+^-H^+^ exchange, which increases RBC pH and promotes osmotic swelling of cells (due to influx of Na^+^ balanced by Cl^−^ uptake via Cl-HCO ^−^ exchange) and serves to increase hemoglobin oxygen affinity (Nikinmaa and Salama, 1998). These changes indicated that the experimental conditions elicited physiological responses to low oxygen early (within 6 h) and that they persisted for the duration of the study (24 h).

### Increased HIF1α protein is an early response to acute hypoxia

Under these conditions, HIF1α protein was higher in brain, ovary, and skeletal muscle from fish exposed to hypoxia compared with normoxic controls by 6 h, and HIF1α protein remained elevated in brain and ovary at 24 h. Although there was a trend toward higher HIF1α protein in liver at both 6 and 24 h, this was not statistically significant due to high variability among individuals (see below). In an earlier study of *F. grandis* using similar experimental approaches, Gonzalez-Rosario (2016) showed that HIF1α protein increased in the same tissues after 24 h exposure of *F. grandis*, and Borowiec et al. (2018) reported higher levels in skeletal muscle after 12 h exposure of *F. heteroclitus*. Law et al. (2006) showed that exposure of grass carp to 0.5 mg O_2_ l^−1^ for 4 or 24 h increased HIF1α protein in liver, and Rissanen et al. (2006) reported that exposure of Crucian carp to 0.7 mg O_2_ l^−1^ for 6 – 48 h led to higher HIF1α in various tissues (liver, heart, gills, and kidney), although the magnitude of the increase differed among tissues and depended upon exposure duration and temperature. More recently, O’Brien et al. (2020) documented increased HIF1α protein in heart of Antarctic icefish, *Notothenia coriiceps*, after 12 h of hypoxia. Taken together, an increase in HIF1α protein in many tissues appears to be part of the early response of several fishes to acute hypoxic exposure.

We found that HIF1α protein levels in *F. grandis* gill are very low and unaffected by hypoxia. This observation contrasts with that reported for Crucian carp, which display substantial levels of HIF1α during normoxia that are further increased by hypoxia (Sollid et al., 2006; Rissanen et al., 2006). The increase during hypoxia (0.7 mg O_2_ l^−1^) in Crucian carp gill depended upon temperature, occurring at 8°C, but not at 18°C or 26°C. Sollid et al. (2006) proposed that HIF1α protein plays a role in the remodeling of gills to increase the surface area for gas exchange during hypoxia (Sollid et al., 2003). Thus, it is possible that the difference in gill HIF1α protein abundance between Crucian carp and *F. grandis* correlates with their capacity for gill remodeling. Although this process has not been examined in *F. grandis*, it was not observed in the closely related *F. heteroclitus* during hypoxic exposures at 21°C (Borowiec et al., 2015).

There was considerable variability in the tissue levels of HIF1α protein among individuals, especially during hypoxia, suggesting that individual fish are differentially impacted by low oxygen. This suggestion is supported by the observation that blood indicators of hypoxic exposure also showed considerable variation among individuals. Fish with higher blood lactate concentrations during hypoxia also had higher levels of HIF1α protein in liver tissue (Spearman’s rho = 0.64, p = 0.006), although this relationship was not observed for other tissues. It is possible that variation in body mass also contributes to individual variation in HIF1α protein levels, as documented for Crucian carp (Sollid et al., 2006). In the current study, gill HIF1α protein levels in hypoxic *F. grandis* were significantly positively related to body mass (Spearman’s rho = 0.69, p = 0.004), opposite of the negative relationship reported for gill HIF1α protein in normoxic and hypoxic Crucian carp (Sollid et al., 2006). Finally, *F. grandis* HIF1α protein levels were not correlated among tissues, in contrast to the positive correlations across several tissues in normoxic Crucian carp (Rissanen et al., 2006). At present, the causes and potential consequences of individual variation in tissue levels of HIF1α protein are largely unknown.

### HIFα mRNAs do not increase during short-term hypoxia in *F. grandis*

Unlike HIF1α protein, HIF1α mRNA levels in tissues of *F. grandis* were unaffected by short-term hypoxic exposure. In other fishes, the effects of hypoxia on tissue levels of HIF1α mRNA vary considerably (see Introduction). In liver, HIF1α mRNA has been reported to increase during short-term exposure (< 24 h) to levels of hypoxia similar to those used here (0.4 – 2.0 mg O_2_ l^−1^) in about half of the species studied, while remaining unchanged in the others (Dataset S1). Even within a species, results can differ between studies using similar experimental conditions. In yellow catfish (*Pelteobagrus fulvidraco*), Wang et al. (2020) reported elevated levels of liver HIF1α mRNA after exposure to 1.14 mg O_2_ l^−1^ for 1 and 3 h, but not 6 h, whereas Pei et al (2021) saw the opposite results, where HIF1α mRNA was unchanged at 1.5 and 4 h of exposure to 0.70 mg O_2_ l^−1^, but higher at 6.5 h. In gill, levels of HIF1α mRNA were unchanged in largemouth bass (*Micropterus salmoides*) and tilapia (*Oreochomis niloticus*) exposed to levels of hypoxia similar to those used here (Yang et al., 2017; Li et al., 2017). On the other hand, Wang et al. (2017) documented higher levels HIF1α mRNA in yellow croaker (*Larimichthys crocea*) at several timepoints between 1 and 24 h, although at higher levels of oxygen than those used here. Although fewer studies have measured HIF1α mRNA in skeletal muscle and ovary, levels of this transcript were generally unaffected by hypoxia (Dataset S1), except that longer exposures (3 – 7 days at 1.70 mg O_2_ l^−1^) led to higher HIF1α mRNA in ovary of the Atlantic croaker (*Micropognias undulates*) (Rahman and Thomas, 2007).

We also measured the levels HIF2α and HIF3α mRNA in tissues of *F. grandis* and found no effects of acute hypoxic exposure. The limited reports on the effects of hypoxia on HIF2α mRNA in fishes reveal variation among tissues and species similar to HIF1α mRNA (Dataset S1). In particular, increases have been documented in gill and liver (although not in all species), but generally not in muscle or ovary, (except for much longer exposures) (Dataset S1). Although HIF3α is least studied form, it is broadly expressed in tissues of a variety of fishes (Townley et al., 2022). This gene was originally described in the grass carp (*Ctenopharyngodon idellus*) as HIF4α (Law et al., 2006), but it was subsequently grouped with HIF3α from other ray-finned fishes (Townley et al., 2022). Law et al. (2006) exposed grass carp to 0.5 mg O_2_ l^−1^, sampled fish at 4 and 96 h, and measured levels of HIF3α mRNA in several tissues by northern blotting. HIF3α mRNA was higher than normoxic controls in all tissues at both durations of hypoxia, except skeletal muscle at 4 h.

While we failed to detect changes in HIF1α, HIF2α or HIF3α mRNA in tissues of *F. grandis*, the accumulated evidence suggests that these transcripts increase in abundance in certain tissues during hypoxic exposure of selected species. The reasons for differences among studies are unknown but likely arise from biological (e.g., species) and technical (e.g., experimental exposures) variation among studies. Also, we cannot exclude the possibility that longer hypoxic exposures cause changes in mRNA levels of HIFα transcripts in *F. grandis*, as proposed for HIF2α in ovary of Atlantic croaker (Rahman and Thomas, 2007). As pointed out by others, however, caution should be exercised when interpreting changes in mRNA levels because it is not certain if the corresponding proteins increase, nor that they activate downstream gene expression (Mandic et al., 2021).

### Implications for the mechanism of HIF1α protein increases during acute hypoxia

The current study showed that short-term exposure to hypoxia led to higher mean levels of HIF1α protein in certain tissues, even when mean mRNA levels were unchanged. These results are similar to those reported for fish cells in culture, early developmental stages, and adult tissues (Soitamo et al., 2001; Law et al., 2006; Rissanen et al., 2006; Sollid et al., 2006; Robertson et al., 2014; Guan et al., 2017). We demonstrated for the first time, to our knowledge, that HIF1α protein levels do not correlate with HIF1α mRNA levels when measured in the same tissues among individual fish exposed to either normoxia or hypoxia. Both lines of evidence argue that new transcription is not required for the initial increase in HIF1α protein levels in tissues of *F. grandis* exposed to hypoxia. Rather, the current results support protein stabilization as the mechanism underlying the increase in HIF1α protein during short-term hypoxia as reported for mammals and fish cells in culture (Kaelin and Ratcliff, 2008; Soitamo et al., 2001). The current study reinforces the importance of measuring HIF1α protein levels, and ultimately target gene expression, in order to better understand the molecular responses of fish to low oxygen.

## List of Symbols and Abbreviations

ANOVA: analysis of variance
ARNT: aryl hydrocarbon nuclear translocator
CN: copy number
Hb: hemoglobin
HIF: hypoxia inducible factor
MCHC: mean corpuscular hemoglobin concentration
MCV: mean corpuscular volume
PHD: prolyl hydroxylase domain
qPCR: quantitative real-time polymerase chain reaction
RBC: red blood cells

## Acknowledgements

We thank Emily Martin for help with red blood cell and hemoglobin measurements and Jenna Hill for assistance with database searching for hypoxia effects on HIFα mRNA in fish.

## Data Availability

Data generated by this study will be deposited in an appropriate public database (e.g., Figshare) upon manuscript acceptance.

